# Predator-like auditory stimuli elicit escape climb in *Aedes aegypti* mosquitoes

**DOI:** 10.64898/2026.09.01.748447

**Authors:** Michael J. Rauscher, Gabriella H. Wolff

## Abstract

The yellow fever mosquito *Aedes aegypti* possesses an acute sense of hearing that mediates a well-described courtship and mating behavior^1^, but the role that mosquito audition plays in other important behavioral contexts has been comparatively less understood. Like many other animals, *Ae. aegypti* exhibit escape responses to threatening visual and wind stimuli^2,3^, the latter of which are sensed via the same antennal mechanosensory organ that mediates hearing. Here, we investigated the possibility that mosquitoes may also use their sense of hearing to detect and respond to the acoustic signature of aerial predators such as dragonflies. In free flight experiments, we characterize a rapid climb maneuver in response to low frequency sounds (180Hz and 300Hz tones) as well as spectrally similar recorded dragonfly audio, distinct from responses observed upon presentation of conspecific wingbeat tones of equivalent acoustic loudness. Presentation of these sounds in tethered flight experiments show wing kinematic modulation comparable to that observed in response to visual loom. Taken together, these findings are congruent with the hypothesis that mosquitoes initiate a rapid climb in response to threatening auditory stimuli from aerial odonate predators. This behavior may be exploited for vector control strategies incorporating audio devices.

**Significance Statement:** Mosquitoes possess sensitive hearing that is used during courtship flights but whose role has not been thoroughly investigated in other behavioral contexts. We demonstrate that low frequency tones (180Hz and 300Hz) as well as spectrally similar recorded dragonfly audio elicit a rapid climb in mosquitoes. Together with tethered flight experiments showing that mosquitos modulate their wingbeats in response to low frequency tones similarly to looming visual stimuli, our work suggests that mosquitoes may use their hearing to detect and avoid aerial predators. Further understanding of this sensory pathway may aid in efforts to control mosquito borne illnesses by informing efforts to disrupt mating, biting, and oviposition behaviors.

## Introduction

Many animals exhibit escape behaviors in response to threatening stimuli, balancing selective pressures to optimize rapidity of response while retaining behavioral flexibility^4^. Well-studied arthropod examples include *Drosophila* takeoff evoked by visual loom, touch-mediated crayfish tailflip^5^, and airflow-evoked escape in cockroaches mediated by the cercal hairs^6^. Threat stimuli in all these examples activate a dedicated channel that bypasses ordinary action selection processes, offering a window into mechanisms of neural control and decision-making. Flying *Aedes aegypti* mosquitoes initiate escape maneuvers prompted by both visual and airflow cues, sensing the latter using the antennal Johnston’s organ. This mechanosensory system also mediates hearing in *Ae aegypti,* sensitive to sounds up to 2000Hz^7^ at distances of up to ten meters^8^, and raises the possibility that sound stimuli may also elicit escape flight.

Prior research has focused on audition’s role in courtship and mating, in which mosquitoes listen for conspecific wingbeat frequencies^1^. Though recent work in smaller *Anopheles* mosquitoes has established a role for hearing in collision avoidance^9^, comparatively little is known about sound-mediated behavior in mosquitoes beyond these conspecific signaling contexts. Here, we characterize a rapid climb response in free-flying mosquitoes elicited by low-frequency sounds (180Hz and 300Hz) and recordings of dragonfly wingbeats containing these frequencies. Equivalently loud tones reflecting female (450Hz) and male (700Hz) conspecifics did not elicit the climb response. In tethered flight, these sounds elicited modulation of wing kinematics comparable to those observed in the escape response to visual loom. Taken together, these data are consistent with the hypothesis that mosquitoes use hearing to detect and avoid aerial predators such as dragonflies.

Fieldwork has demonstrated that male *Aedes diantaeus* mosquitoes exhibit a negative phonotaxis in response to low frequency sounds in the range of 100-250Hz^10^. While the ecological relevance of these stimuli has remained unresolved, we hypothesize that sounds in this frequency range may represent the acoustic signature of aerial predators such as dragonflies and damselflies. Compared with other Diptera, mosquitoes are not nimble fliers, exhibiting slower takeoffs and body rotation rates compared with similarly-sized Brachyceran flies such as house flies and bottle flies^11^. Mosquitoes also possess wing motor systems that (due to optimization for their acoustic signaling role) are categorically less efficient in the production of flight forces than those of other flies^12^. Nevertheless dragonflies paired with individual potential prey animals in an enclosed arena demonstrated poorer capture rates pursuing *Aedes* mosquitoes than against *Drosophila,* whose small size should present a more difficult target^13^. While sensory aspects of dragonfly prey perception may explain this phenomenon, here we explore the possibility that mosquitoes make use of audition to perceive a predation attempt and react evasively.

## Results

### Low-frequency sound stimuli evoke rapid climb in free-flying mosquitoes

To investigate whether mosquitoes use audition to mediate aerial escapes, we began by surveying their responses to a variety of sound stimuli in flight. We developed an automated system to play an experimental sound and record high speed video when a mosquito flew into a defined volume (Figure 1A). In addition to a no-stimulus control to provide a baseline of comparison (Figure 1Bi), we presented the animals with four experimental tones: a 180Hz tone (Figure 1Bii) near the center of the previously identified range of putatively aversive stimuli^10^, a 300Hz tone (Figure 1Biii) within the receptive range of the mosquitoes’ auditory system but without explicitly hypothesized ecological relevance, and 450Hz and 700Hz tones representing the respective wingbeat frequencies of female and male *Ae. aegypti* mosquitoes that may elicit conspecific-related behaviors (Figure 1Biv-v).

**Figure 1.**
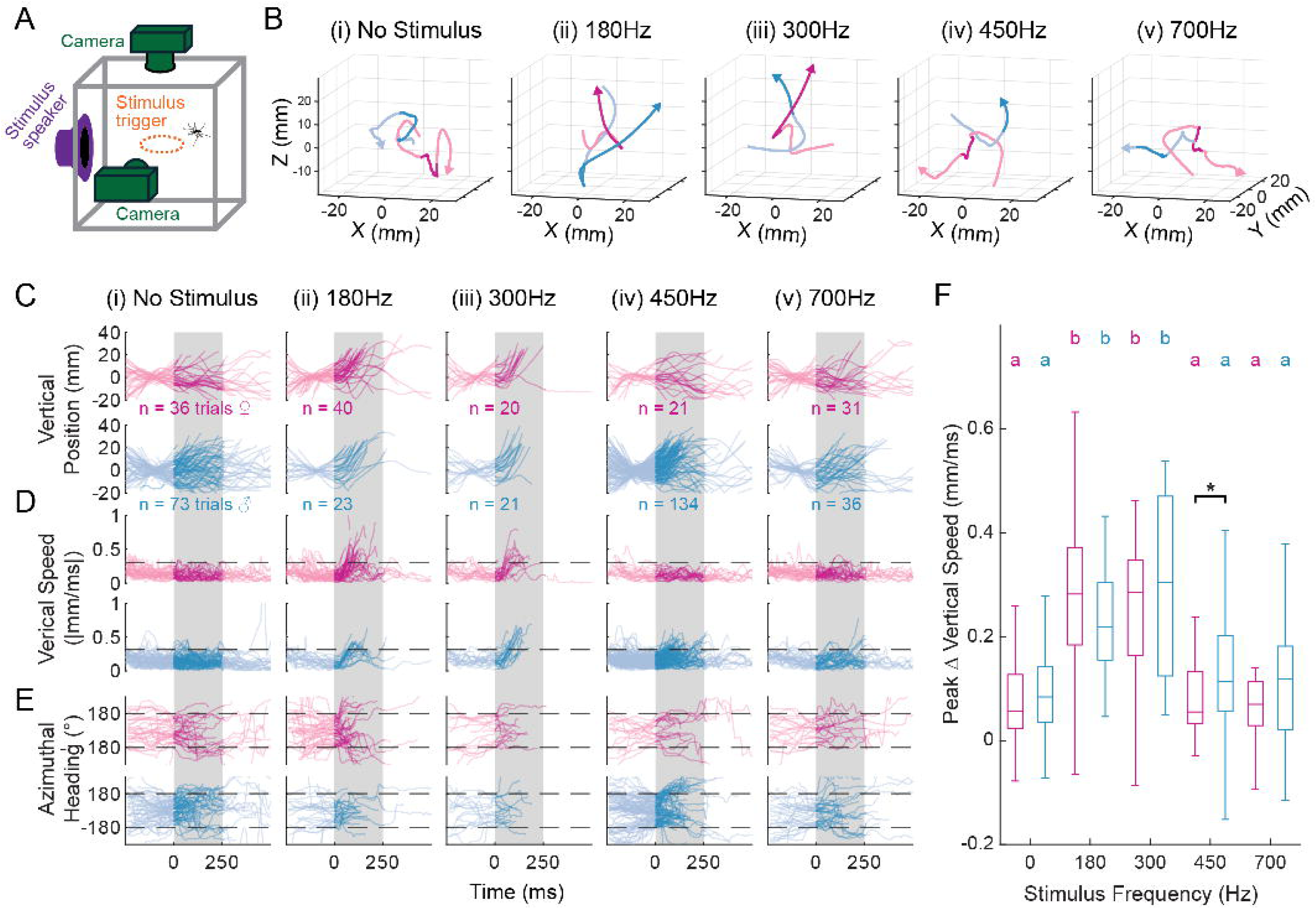
Male and female mosquitoes initiate rapid climb in response to low frequency sounds. **(A)** Schematic view of automated freeflight behavioral arena, showing vertical and horizontal aspect cameras, stimulus loudspeaker and central region of real-time behavioral monitoring. **(B)** Representative trajectories for (i) control trials with no stimulus, (ii) 180Hz and (iii) 300Hz low frequency stimuli alongside (iv) 450Hz and (v) 700Hz stimuli within the wingbeat frequency range of female and male *Aedes aegypti* respectively. **(C)** Vertical position of each side-aspect trajectory over time, zeroed on each trajectory’s pre-trial baseline. Upper axes show female trajectories in magenta, with males shown in blue in lower axes. **(D)** Same as C, but showing speed (magnitude of velocity vector) in the vertical aspect. Dashed line shows three standard deviations from the mean. **(E)** Direction of travel over time, with zero facing the stimulus speaker. **(F)** Comparison of peak vertical speed from each trajectory (shown in B) between sexes and stimulus classes. Magenta and blue boxplots summarize female and male data respectively, with colored letters summarizing statistically distinct subsets within each sex (p< 0.05 One-way ANOVA familywise test with Tukey-Kramer *post-hoc* test). Black asterisk marks stimuli with statistically distinguishable differences between the two sexes (* = p < 0.05, two sample t-test), showing that males exhibit elevated vertical speed in response to the 450Hz female tone.

We measured each animal’s vertical position (Figure 1C) and velocity (Figure 1D) as well as their azimuthal heading (Figure 1E). In the majority of observed trajectories, stimulation with the 180Hz and 300Hz tones resulted in the initiation of a rapid climb maneuver (Figure 1C) with an accompanying significant increase in vertical flight speed (Figure 1D,F). If the observed climb behavior represents an escape response, we hypothesize that it should be distinct from responses to conspecific stimuli. In agreement with this hypothesis, we did not observe comparable increases in vertical (Figure 1F) or translational (Figure S1A) flight speed upon presentation of the 450Hz and 700Hz tones of comparable sound pressure level (representing conspecific wingbeat frequencies). These findings suggest that the animals were not merely responding to the acoustic loudness of the lower frequency stimuli *per se,* but also to the spectral content of the stimulus. Though mosquitoes frequently changed their direction of travel following stimulus presentation (Figure 1E), we did not observe a discernable relationship between the mosquito’s initial heading (relative to the stimulus) and its ultimate direction of travel within the following 500ms (Figure S1B). These results are congruent with prior work quantifying escape flight parameters in response to a simulated swatting attack, which emphasized that baseline levels of stochasticity in the flight path contributes to escape success in addition to active escape maneuvering^2^.

Reflective of their unique acoustic signaling system, male and female mosquitoes exhibit considerable sexual dimorphism in their body size, antennal morphology, and auditory sensitivity^14–16^. If low frequency tones signal threat, we would predict that male and females would respond similarly to them, as both are subject to predation risk. Indeed we did not observe significant differences in vertical flight speed (Figure 1F), translational flight speed (Figure S1A), or heading (Figure S1B) between males and females in their responses to the low frequency 180Hz and 300Hz stimuli. Whereas neither males nor females responded to the 700Hz tone (produced by male wingbeats), we observed that male mosquitoes responded to the 450Hz tone (produced by female wingbeats) by increasing their vertical flight speed significantly higher than females (Figure 1D). Unlike with the low frequency stimuli, however, this increase did not exceed the range of flight speeds observed under the no-stimulus condition (Figure 1F). Similarly, males presented with the 450Hz female tone tended to end their trajectories on a different heading than their pre-stimulus baseline, potentially reflective of searching behavior (Figure S1B). Taken together, our data show that both males and females respond similarly to low frequency sound stimuli, and that male mosquitoes respond to female wingbeat tones in a qualitatively different manner that is consistent with our prior understanding of the role of audition in courtship.

### Tethered mosquitoes increase wing motor output in response to low frequency sound

To better assess potential sex differences and more definitively characterize the biomechanical outputs mediating this behavior, we presented the same sound stimuli to mosquitoes in tethered flight (Figure 2A). For a mosquito to accelerate and increase its altitude, as observed, it must increase its wing thrust, which it can accomplish by increasing the total wingstroke amplitude (Θ_L_+Θ_R_) and/or the wingstroke frequency. To measure these, we extracted the stroke amplitude from the high-speed video data using Flyalzyer^17^, a MATLAB-based machine vision tool (Figure 2B-C), and the stroke frequency using one of several machine vision methods based around the Fast Fourier Transformation (Figure 2C, see Methods). In tethered flight, steering effort can also be estimated by estimating wing yaw as the difference between the two wing amplitudes (Θ_L_+Θ_R_, Figure 2B-C), in concert with the head yaw angle relative to the body axis (Figure 2B).

**Figure 2.**
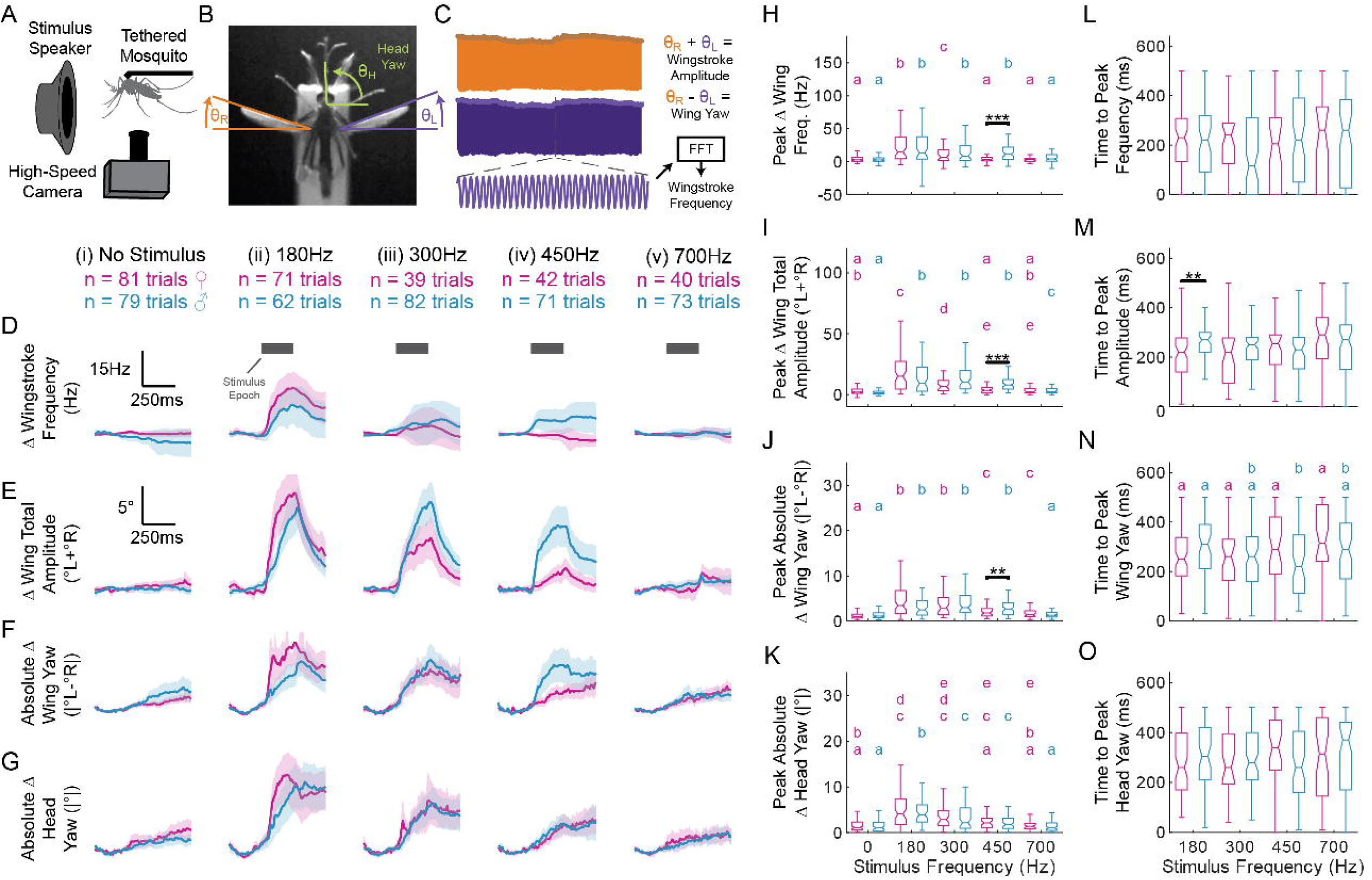
Female mosquitoes respond more quickly to low frequency auditory stimuli than males. **(A)** Schematic view of tethered flight behavioral rig, showing tethered mosquito, stimulus loudspeaker, and ventral aspect high-speed camera. **(B)** Measured kinematic parameters, including wing leading edge and head angular position in the yaw aspect. **(C)** Computed kinematic parameters, illustrating total total wingstroke amplitude (Θ_L_+Θ_R_) as the sum of the downstroke envelopes of the two wings, wing yaw as their difference (Θ_L_+Θ_R_), and wingbeat frequency computed via the fast fourier transform over a sliding window of high speed video frames. **(D)** Mean (dark lines) and 95% confidence interval (shaded regions) for wing frequency deviation from baseline. **(E)** Same as D, but showing total amplitude (Θ_L_+Θ_R_) change from baseline. **(F and G)** same as D and E, but showing respectively the absolute value of the wing yaw (Θ_L_-Θ_R_) and head yaw relative to baseline. **(H-K)** Comparisons of peak deviation from baseline for each kinematic parameter during stimulus epoch. Colored letters denote subsets resulting from Kruskal-Wallis omnibus test with Conover-Iman *post-hoc* test within each sex across stimulus conditions, whereas asterisks denote significant differences between sexes within a stimulus condition (** = p< 0.01, *** = p< 0.001, Wilcoxon rank sum test). **(L-O)** same as H-K, except showing time to peak response level for all tones. Female wing total amplitude response (Θ_L_+Θ_R_, M) is significantly faster than males for 180Hz stimulus.

Both wingstroke frequency (Figure 2D,H,L) and wingstroke amplitude (Figure 2E,I,M) increased following presentation of the 180Hz and 300Hz stimuli, with wing amplitude (Θ_L_+Θ_R_) modulation following a faster time course in females versus males for the 180Hz stimulus (Figure 2Eii,M). Wing yaw (Θ_L_+Θ_R_, Figure 2F,J,N) and head yaw (Fig 2G,K,O) estimates largely resembled those observed for wing amplitude (Θ_L_+Θ_R_). Congruent with findings from the free flight experiments, male responses to the 450Hz female wingbeat tone stimulus were of comparable amplitude but qualitatively different from responses to the 180Hz and 300Hz stimuli, with wing frequency remaining elevated following stimulus presentation rather than returning towards baseline (Figure 2Div). These findings broadly agree with our findings in the free flight experiments, and raise the possibility of more finely-grained differences between female and male mosquitoes in their response to low frequency sound than are evident in the free flight trajectories. It is unclear if the faster response of females to low-frequency tones observed in the tethered flight experiments translate consequentially to free flight kinematics.

### Auditory-evoked changes in wing and head kinematics resemble responses to visual loom

If the described auditory-evoked response indeed reflects an escape behavior as we hypothesize, then we expect it should be similar to escape responses evoked by other sensory modalities. Many animals (including *Ae. aegypti*) demonstrate escape behaviors elicited by visual expansion or “looming” stimuli^18,3,2,9^. To elicit this response, we presented the mosquitoes with an expanding square stimulus using a modular LED display^19^. As with the 180Hz auditory stimulus, both male and female mosquitoes responded to visual looms by increasing their wingstroke frequency (Figure 3A,C,D), total wingstroke amplitude (Θ_L_+Θ_R_, Figure 3B,E,F), wing yaw (Θ_L_-Θ_R_, Figure S2A,C), and head yaw (Figure S2B,D). These findings are consistent with the interpretation that the auditory-evoked responses reflect an escape response, as they are comparable in time course and magnitude to those evoked by visual expansion.

**Figure 3.**
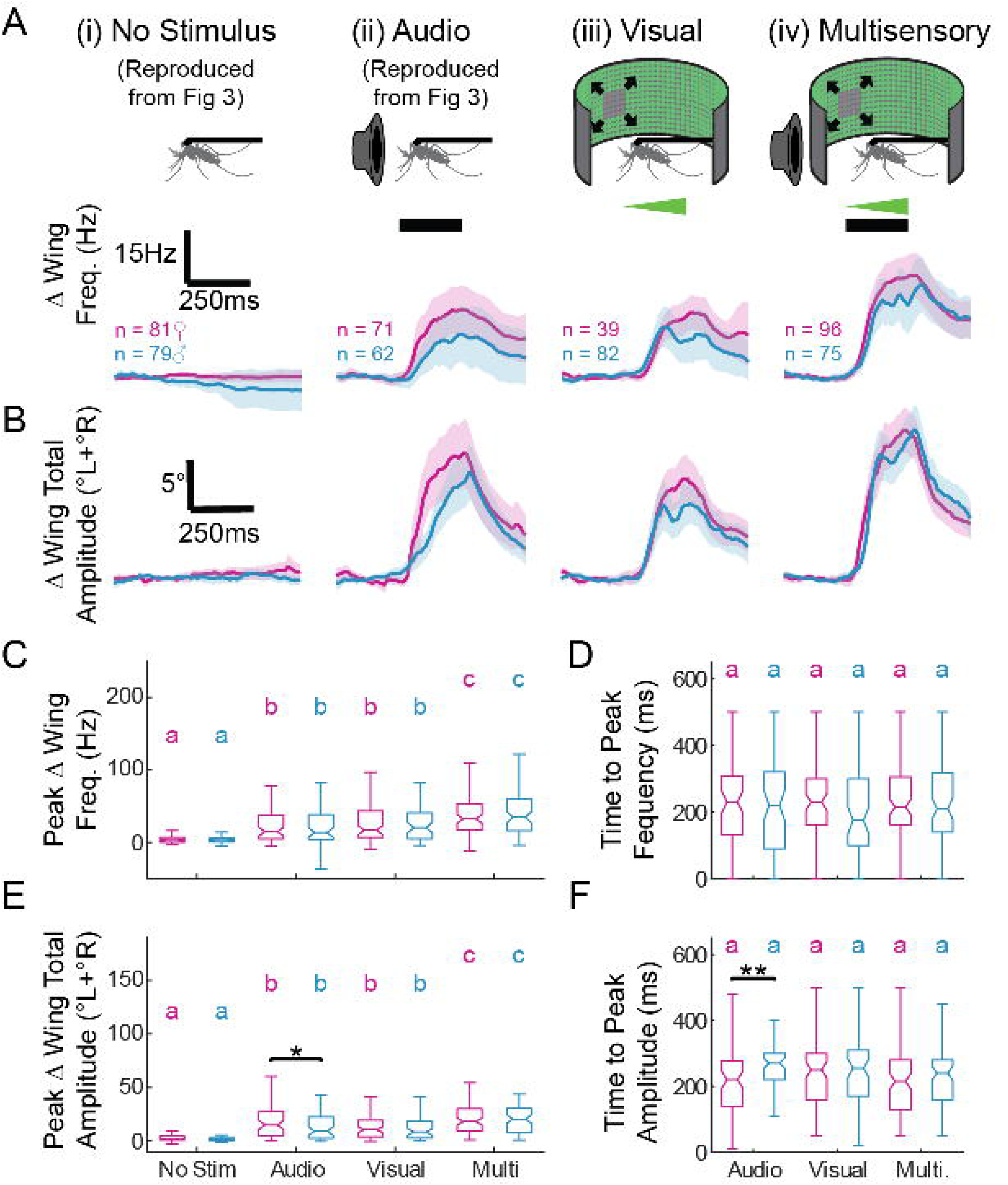
Auditory-evoked response is similar to visual loom response. **(A)** Wingbeat frequency deviation from pre-trial baseline over time, with schematic view illustrating each experimental condition: no stimulus (i) and 180Hz audio stimulus only (ii) reproduced from Figure 3 as well as (iii) presentation of an expanding square looming stimulus and (iv) concurrent presentation of both the looming and 180Hz auditory stimulus. **(B)** Same as (A), but instead showing deviation from baseline for total wingstroke amplitude (Θ_L_+Θ_R_). **(C)** Comparison across sexes and conditions for wing frequency response (colored letters reflect outcome of Kruskal-Wallis omnibus test with Conover-Iman *post-hoc* test within each condition, * = p< 0.05 Wilcoxon rank sum test pairwise between sexes), showing comparable response magnitudes for auditory and visual conditions. Response magnitude for multisensory condition exceeds those observed under unisensory conditions. **(D)** Same as (C), but for time to peak response across conditions, showing males and females responding at equivalent timescales under the visual and multisensory conditions (** = p< 0.01 Wilcoxon rank sum test). **(E-F).** Same as C-D, but for total wing amplitude (Θ_L_+Θ_R_) response.

Our findings also imply that sex-specific differences in the response to auditory stimuli reflect differences in auditory sensory processing, rather than originating from differences in biomechanical production of flight forces. In contrast with the auditory responses, male and female mosquitoes did not differ in their time to peak wing amplitude response when presented with the visual stimulus (Figure 3D). Additionally, while females responded to the looming visual stimulus more slowly in the wing yaw (Θ_L_-Θ_R_, Figure S2D) and head yaw (Figure S2F) aspects, they did not do so to a degree statistically discernable from the range of responses observed in males. Taken together with the free flight data (Figure 1F), in which auditory stimuli evoked similar changes in vertical flight speed, these findings imply comparable ability to produce the observed flight forces between males and females. Further work is required to explore whether the faster female response to low frequency tones reflects a behavioral significance.

### Auditory and visual inputs do not assert exclusive control over escape flight

Escape systems used during flight face the additional challenge of integrating seamlessly with the complex sensorimotor control architecture keeping the animal in a controllable attitude with its flight forces in balance. Smaller night-flying *Anopheles* mosquitoes have been shown to integrate looming visual cues with antennal mechanosensation to guide escape maneuvers^20^. In *Drosophila,* these same stimuli evoke rapid changes in heading known as body saccades, released by a dedicated population of four neurons in the central brain^21^.

Saccades appear to proceed “ballistically” after being evoked by visual stimuli, but their dynamics are continuously modulated by inertial sensory feedback, which controls the duration of the turn^22^. In quiescent *Ae. aegypti* as well as in *Drosophila,* responses to visual expansion evoke escape takeoffs^3,18^, which in the latter are known to be mediated by a giant fiber descending pathway. Antennal mechanosensory inputs also converge upon this same pathway, mediating escape takeoff in response to wind stimuli as well^23^. To assess whether escape responses mediated by audition are integrated with those evoked by vision, or whether they vie for control of a common command pathway, we presented both the 180Hz auditory stimulus and the looming visual stimulus concurrently (Fig 3A*iv*,B*iv).* We observed categorically larger wing frequency responses (Figure 3C) and total wing amplitude responses (Θ_L_+Θ_R_, Figure 3E) than for either stimulus independently. Both kinematic parameters were comparable in their time to peak onset for multisensory as for visual stimuli (Figure 3D,F). Similar results were observed in wing yaw (Θ_L_-Θ_R_, Figure S2A*iv*) and head yaw (Figure S2B*iv*), with unisensory and multisensory responses having comparable amplitude (Figure S2C,E) and time to peak response (Figure S2D,F). Taken together, these findings suggest a non-exclusivity of descending control over escape flight by each sense modality, as the categorical increase in the wing kinematic responses and the flattening of sex-specific differences under multisensory conditions would not be predicted under a “winner-takes-all” control paradigm. Similarly, these results demonstrate that the unisensory responses do not saturate the operating range of the wing motor system.

### Recorded dragonfly audio elicits similar responses to those evoked by low frequency tones

Previous work has established that mosquitoes make extensive use of both visual and airflow cues to anticipate and avoid swatting, capture, or collision risks^2,20^. In these prior studies, the visual and airflow cues from a mechanical swatting device provided a clear experimental analog to the swatting attempts made by large vertebrate hosts. Here, we show that acoustic stimuli mediate a similar behavior in a frequency-dependent manner, suggesting that escape flight initiation is not simply a function of sudden global changes in particle velocity or ambient sound pressure levels. While these data leave the ecological significance of these salient sound stimuli incompletely resolved, one attractive hypothesis is that mosquitoes react evasively in response to the acoustic signature of aerial predators such as dragonflies.

To investigate the plausibility of this hypothesis, we recorded takeoff audio from a dragonfly (*Epitheca canis)* to analyze its spectral content (Fig 4A-B) and observe how mosquitoes respond to it in free flight (Fig 4C-E). Though this dragonfly is not found in the home range of *Ae. aegypti,* there is considerable similarity in the basic biomechanics and neural control of flight across dragonfly lineages^24,25^. Dragonflies maintain independent motor control over all four wings and can adopt different wing “gaits” during takeoff and active pursuit^25^ which contribute to strong harmonics in their acoustic profile. In our example recording, we observed strong peaks corresponding to the first, second and third harmonics of the wingbeat frequency, within the putatively-aversive range of low frequency stimuli suggested by the pure tone experiments (Figure 4B).

**Figure 4.**
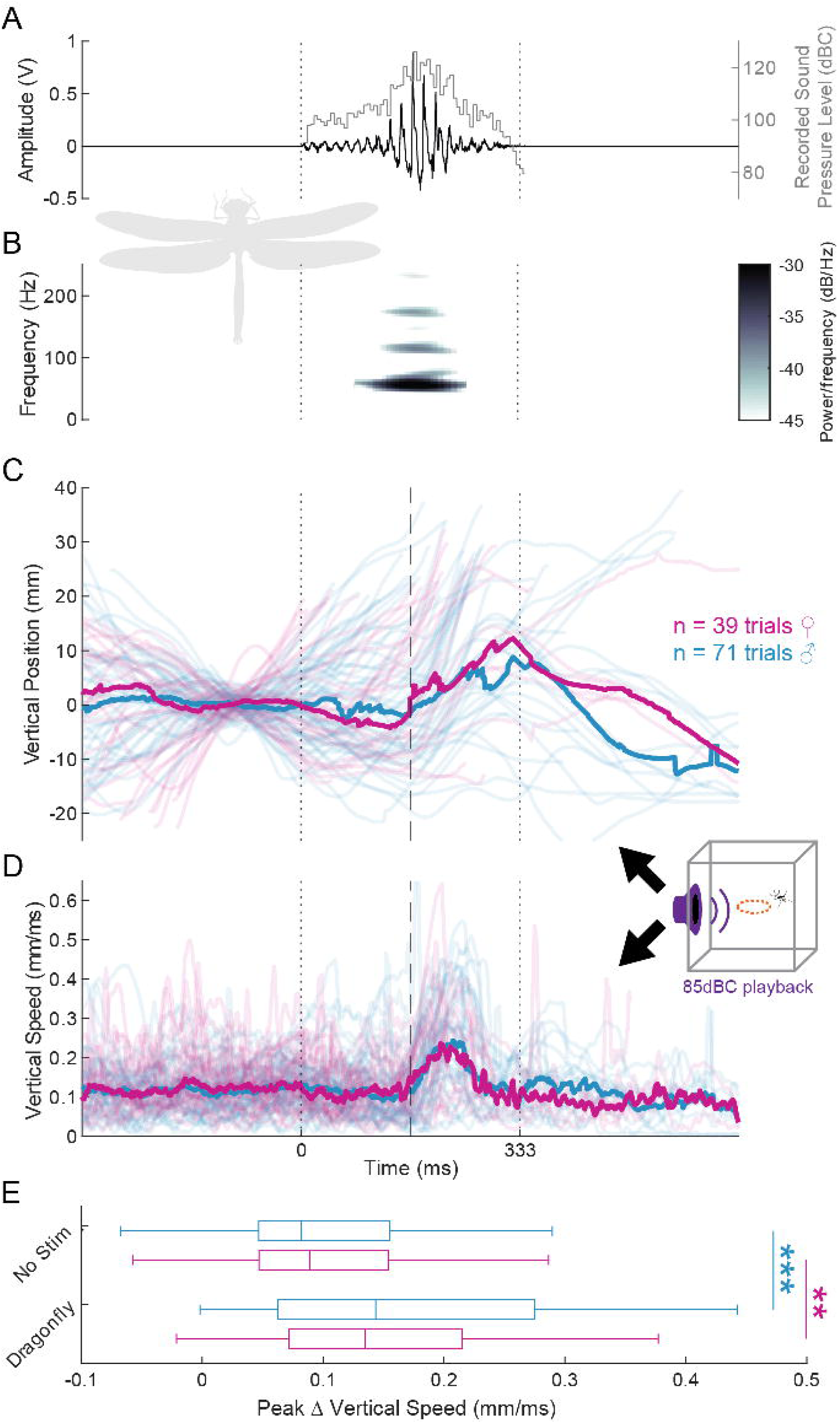
Recorded dragonfly takeoff audio elicits climb response in free flight. **(A)** Voltage over time showing waveform of recorded dragonfly (*Epitheca canis)* audio. Light gray: sound pressure level over time. **(B)** Spectrogram of audio in (A), showing fundamental wingbeat frequency of approximately 60Hz, with strong harmonics evident within the 100-300Hz range. **(C)** Side-aspect trajectories of male (blue) and female (magenta) mosquitoes responding to playback of the dragonfly audio in the freeflight arena, with peak loudness calibrated to 85dBC at the center of the arena. Thick lines represent average response, with apparent discontinuities reflective of animals entering or exiting the field of view of the camera. Dashed line corresponds to the loudest point in the stimulus, eliciting climb response shortly thereafter. **(D)** Same as (C), but showing the vertical speed of each animal, increasing during the climb maneuver and returning to baseline as the stimulus amplitude decreases. **(E)** Comparison of peak vertical speed deviation from baseline against no stimulus control (reproduced from Figure 1, with pre-trial baseline estimate adjusted to match longer dragonfly stimulus), showing significant increase in vertical speed in both sexes (** = p< 0.01, *** = p<0.001, two sample t-test).

Playback of this sound in the free-flight arena elicited stereotyped increases in altitude (Fig 4C) and significant increases in flight speed (Fig 4D-E) similar to those observed in response to the low-frequency pure tone stimuli. These increases in flight parameters corresponded with the loudest part of the stimulus, where the dragonfly initiated its takeoff. In the subsequent part of the recording, where the dragonfly moved away from the microphone and stimulus amplitude diminished, the mosquitoes’ flight speed returned towards baseline for animals remaining in the field of view of the camera (Figure 4D). These data support the role of low-frequency tones as potential signaling cues for aerial predators.

## Discussion

Here, we have characterized an auditory-mediated rapid climb behavior in flying *Ae. aegypti* mosquitoes, and shown that it plausibly reflects an escape response to the acoustic signature of aerial odonate predators. This corroborates prior characterization of an escape climb in response to startling airflow cues in smaller *Anopheles* mosquitoes^26^ and may resemble acoustic escape behaviors exhibited by noctuid moths^27^ and mantises^28^ in response to bat ultrasound. These animals engage in stereotyped dive maneuvers that camouflage their motion signature or cause the bat to lose track of them. The escape climb observed in mosquitoes might reflect a similarly appropriate response to dragonfly predators, which typically attack from below^29,30^ and as visually-guided predators are unlikely to lose track of an evading prey animal. Engaging all available flight motor output may instead reflect a last-moment feint, giving the mosquito the best chance of altering its trajectory outside of the dragonfly’s capture envelope once it is too far along in its terminal guidance to change course any further.

As with other aspects of mosquito audition, the salience of the 100-300Hz range is significant within the context of conspecific courtship behavior, as it contains intermodulation distortion products produced by the interaction of the male and female wingbeats. During courtship flights, *Ae. aegyptes* (like other mosquitoes) modulate their wingbeat frequencies in synchrony to converge on a shared harmonic^7^. Behavioral evidence shows that the ability of a focal male to produce this convergence correlates with mating success in *Ae. aegypti*^31^ and research in other mosquito species suggests that convergence may emerge as a consequence of wingbeat frequency modulation aimed at generating low frequency intermodulation distortion tones optimal for sound localization^32,33^. We therefore cannot discount that our observed behavior reflects an artificial engagement of courtship-related conspecific signaling mechanisms outside of their normal behavioral context. We consider this unlikely however, as playback of the low-frequency experimental tones resulted in statistically indistinguishable free flight outcomes for both males and females, without clear evidence of attempted sound source localization, and qualitatively distinct from the response to conspecific flight tones in both free and tethered flight. We observed the same vertical climb in response to recorded dragonfly audio, which is a predator cue with a distinct spectral signature from any conspecific signal. Additionally, the climb behavior corroborates the negative phonotaxis observed in prior work as well as the escape climbs reported in response to visual loom and wind stimuli^2,26^.

We consider it likely that the described behavior depends principally upon antennal hearing, though it cannot be discounted that other mechanosensory organs apart from the antennae may play a role. Signal transduction in the Johnston’s organ is accomplished via specialized complexes of stretch-sensitive cells known as scolopidia, genetically identifiable in both males and females via the expression of a common TRPV channel necessary for hearing-mediated courtship behavior^34^. Scolipidial cells are also present in chordotonal organs in all of the mosquito’s appendages including the legs, wings, and halteres (reviewed ^35,36^).

Traditionally considered solely as proprioceptors, recent work in *Drosophila* suggests that they may contribute to the detection of exteroceptive vibrations beyond those mediated by antennal hearing^37^. Future work will be necessary to pinpoint the location of auditory sensors that transduce low-frequency tone stimuli.

More broadly, the public health challenges posed by blood-feeding mosquitoes motivate study of all aspects of their basic biology. Better understanding of sensory mechanisms mediating acoustically-driven behavior during flight may point to novel modes of disrupting other behaviors including mating, biting, and oviposition^38^. Similarly, our work motivates closer attention to behavioral strategies employed by mosquitoes in interaction with predators, as these may further aid in vector control efforts by giving insight to the ways that mosquitoes interact with the built environment and facilitate design of better traps, lures, and assays.

## Methods

| REAGENT OR RESOURCE | SOURCE | IDENTIFIER |
| --- | --- | --- |
| <u>Deposited Data</u> |  |  |
| Analysis Code | Github.com | https://github.com/michaelrauscher/ |

| <u>Experimental Models: Organisms/Strains</u> |  |  |
| --- | --- | --- |
| *Aedes aegypti* | The following reagent was obtained through BEI Resources, NIAID, NIH: *Aedes aegypti*, Strain LVP-IB12, Eggs, MRA-735, contributed by David W. Severson | Strain LVP-IB12, Eggs, MRA-735 |
| <u>Software and Algorithms</u> |  |  |
| MATLAB 2025B | Mathworks Ltd. | http://www.mathworks.com/products/matlab/; RRID:SCR_001622 |
| <u>Other</u> |  |  |
| TS4 and IL5 High speed cameras | Fastec Imaging | https://www.fastecimaging.com/ |
| Nidaq USB-6212 and USB-6343 | National Instruments | https://www.ni.com/ |
| Nobsound G2 Pro Amplifier | Douk Audio | https://www.doukaudio.com/ |
| Dickinson-Reiser "Panels" LED Arena | IO Rodeo | https://iorodeo.com/ |

### Experimental model and study participant details

#### Animals

We established a colony of wildtype yellow fever mosquitoes using eggs obtained through BEI Resources, NIAID, NIH: *Aedes aegypti* (Strain LVP-IB12, Eggs, MRA-735, contributed by David W. Severson). Mosquitoes were reared in an incubator at 70C and 27% relative humidity on a 16 hour light period / 8 hour dark period rotation. Animals were kept for seven days post-eclosion before use in behavioral experiments. Larvae were fed commercially available fish food, with adults feeding on a 10% sucrose solution. Animals not retained for experiments were transferred to a reproductive flight cage in the same incubator for blood feeding and egg collection.

### Method details

#### Tethered flight behavioral experiments

Mosquitoes were cold-anesthetized and transferred to a custom peltier-cooled tethering platform for positioning, then subsequently attached to custom 3D-printed tethering sticks by applying a small drop of UV-curing dental cement to the tethering stick and gently sliding the anesthetized mosquito forward until its anterior pronotum contacted the glue drop. Glue was then cured for five to ten seconds under a UV flashlight.

Mosquitoes were then transferred to a 3D hand positioner in the tethered flight behavioral arena and aligned into the field of view and focal plane of a Fastec IL5 high speed camera recording the animal from the ventral aspect at 2500FPS. A subset of animals were recorded instead at 5000FPS for validation of wing kinematic analysis methods (see below *Tethered flight head and wing kinematic analysis)*.

#### Free flight videography and event-triggered recording

We adapted an off-the-shelf 30×30×30cm mesh insectory into an automated free flight behavioral arena capable of detecting a mosquito within a defined space and triggering a sound stimulus and high speed video recording (Figure 1A). A pair of synchronized Fastec TS4 high speed cameras recording at 2000FPS from the overhead and lateral aspects enabled 2D tracking of individuals within each aspect and 3D trajectory reconstruction for many individuals moving within the overlapping fields of view of both cameras. In all trials, we designated the stimulus trigger zone as a cylindrical volume in the center of the arena, with the sugar feeding cup located several centimeters beneath it to attract mosquitoes into the field of view. To validate our set-up and provide a baseline of comparison for flight kinematics, we collected a number of no-stimulus control videos by setting the system to trigger recording without accompanying sound (Figure 1B-Ei). Typically, we recorded with both males and females present in the flight cage, with post-hoc identification of sex possible in the majority of trials, though some recording sessions were conducted with single-sex cages.

The HDMI monitor output from each camera, updating at 30FPS, was sent to a USB video capture dongle which fed the images into a custom MATLAB control and stimulation program *via* the Image Acquisition Toolbox, facilitating real-time detection of mosquitoes in the arena. Prior to a recording session, the amplifier was adjusted manually to achieve the intended sound pressure level. Because temperature and humidity have been shown to influence physiological and behavioral responses to sound stimuli, the arena was placed within the mosquitoes’ home incubator at a constant temperature of 27C and 70% relative humidity.

#### Auditory Stimuli

Experimental pure tone stimuli were sine waves of 250ms duration calibrated to a sound pressure level (SPL) of 80dBC for free flight experiments and 60dBC for tethered flight experiments, as measured using a hand meter placed in the center of the arena, ensuring each sound was approximately 20dBC louder than the ambient background in the incubator and tethered flight behavioral arena respectively. All stimuli fell within the range of uniform input-response for the dBC intensity adjustment scale, permitting direct comparison of sound pressure levels across frequencies.

Dragonfly audio was recorded with a small lavalier microphone placed ∼4cm from the animal as it perched on the side of a mesh flight cage, digitized using a PC sound card. SPL estimates were made by placing a speaker in the perch location and playing a 1KHz reference tone into the same recording system, measured independently with the hand meter. The reference tone and value were then used as calibration factors for the *splmeter* system object in MATLAB to estimate the acoustic loudness of the dragonfly waveform. Playback of this waveform in the free flight arena was adjusted to 85dB, comparable to the value chosen for the pure tone stimuli.

Auditory stimuli were presented to the animals in free flight *via* an 8*Ω*, 1W loudspeaker 100mm in diameter. Tethered flight experiments used one of a pair of 4*Ω,* 3W loudspeakers 50mm in diameter, each placed 45° in azimuth to the left and right of the body axis. Signals were driven via a DAQ (Nidaq USB-6212 or USB-6343, National Instruments, Austin, TX, USA) and sent to an audio amplifier in line with the speakers (Nobsound G2Pro, Douk Audio, Hong Kong, China)

#### Visual Stimulus

Expanding visual loom stimuli were presented via a Dickinson-Reiser cylindrical LED arena^19^ approximately 380mm (96 pixels) in circumference and 140mm (24 pixels) tall, with each pixel representing a 3.75° region of the visual field. Stimuli took the form of a single pixel dot, centered approximately 45° in azimuth to the left or right of the body axis, that grew linearly to a 29×29 pixel square (108.75°) over the course of 250ms.

### Quantification and statistical analysis

Conover-Iman familywise tests were performed in the open source R statistical programming environment. All other data annotation and analysis tasks were performed using MATLAB (The Mathworks, USA).

#### Free flight trajectory tracking and annotation

A custom MATLAB program facilitated offline tracking and annotation of mosquitoes in free flight videos. Average frames were computed for each video to facilitate tracking via a guided background-subtraction method, in which the user would click on a mosquito around which the computer would define a small region of interest (of a size set by the user but typically 16×16 or 32×32 pixels), advance to the next frame, perform a binary thresholding operation using the *imbinarize* function, and locate the animal as the centroid of all active pixels within the defined region of interest, which would in turn center the region of interest for tracking in the subsequent frame.

Manual tracking was performed where this algorithm performed poorly, which were typically segments of video where the trajectories of multiple mosquitoes overlapped and the computer locked on to the wrong animal. The tracking interface facilitated assignment of trajectories from each camera to a common individual, also including fields for sex-identification and qualitative notes about each trajectory (take-off, landing, collision, mated pairs flying together, etc). As our analyses considered time series from each camera separately and the overlapping field of view of the two cameras was restricted to the center of the sensor area, minimizing nonlinear distortion from lens effects, we used a simple linear correction factor to convert image coordinates to real-world units.

#### Free flight trajectory analysis

Absolute vertical and translational speed (defined as the magnitude component of the animal’s velocity vector in the side or top aspect camera respectively) were estimated first by smoothing each position time series using a 15-sample moving average filter, calculating the distance between successive points in the time-series using the euclidean distance formula, and then computing the numeric differential. For each trajectory, peak vertical or translational speed deviation in each aspect was defined as the difference between the maximum value observed during the stimulus epoch and the average speed of all time points in the 250ms pre-stimulus epoch. Azimuthal heading was computed by using the inverse tangent function to measure the angle change between successive frames of the smoothed top-down position time series. The unwrapped azimuth time series was then adjusted such that the value at stimulus onset was wrapped to the interval [-180°:180°].

#### Tethered flight head and wing kinematic analysis

In the majority of experimental trials, wing frequency estimates were calculated by creating a time series of the summed pixel intensity of each image frame. Time-series were Z-normalized to remove low frequency “DC-offset” components in the subsequently calculated spectrograms. Spectrograms were computed with 10ms non-overlapping windows, and the time-frequency ridge (corresponding to the wingbeat frequency) extracted *via* the *tfridge* function in MATLAB. Ridges were visualized alongside their originating spectrograms using a custom proofreading GUI tool to ensure the function had not erroneously locked onto harmonics or aliases of the signal, and such errors corrected by restricting the frequency range over which the spectrogram was calculated to values appropriate for each animal. For a subset of ridges, clearer spectral separation from the background was apparent in the proofreading tool for the first harmonic of the wingbeat frequency signal rather than at the fundamental frequency, and so the time-frequency ridge following the first harmonic (divided by two) was used to estimate the wingbeat frequency for these trials.

Wingstroke amplitude estimates for each wing were made by producing a series of maximum-intensity projections from all frames within non-overlapping 10ms windows of the video (matching the windowing of the wing frequency spectrograms to create aligned, downsampled kinematic time series at 100Hz) to visualize the stroke envelope of the wings. The leading edge of the envelope (corresponding to the wingstroke amplitude) was then extracted using a custom MATLAB program called Flyalyzer. “Wingstroke amplitude” as used throughout the manuscript refers to total wingstroke amplitude, defined as the sum of stroke amplitudes of each wing (Θ_L_+Θ_R_), whereas “wing yaw” refers to the difference of the left and right wingstroke (Θ_L_-Θ_R_). The head yaw angle with respect to the body axis was also estimated in Flyalyzer by tracking the base of the proboscis relative to the neck or in some cases the base of the antennae.

As noted above, a subset of animals were recorded at 5000FPS, this enabled us to track the complete wingstroke with higher temporal resolution, allowing us to validate the methods detailed precedingly for stroke frequency and amplitude estimation. All 139 videos (encompassing 2425689 frames and 48513 FFT windows) achieved R^2^ values in excess of 0.9 comparing wing frequency ridges between the two methods, with 90% of videos exceeding an R^2^ of 0.975.

## Acknowledgements

We would like to thank Kim Thompson for her care of the mosquito colony, as well as Haaris Khatri, Drishti Mangani, Frances Miller, Skylar Monjure, Apple Patel, and Lauren Roskuzka for help with data collection, annotation, and analysis. We would also like to thank Shivansh Dave, Alexandra Gurgis, Jessica Hearn, Kristianna Lea, Amy Streets, Alexandra Yargar, and Nicholas Kathman for helpful commentary and feedback, alongside Andrew Dacks, Jessica Fox, and other members of their respective laboratory groups. Additionally, we would like to thank Jaramillo Hector Sergio for the outline of the *Aedes aegypti* mosquito used in Figures 2 and 3 as well as Gareth Monger for the outline of the dragonfly in Figure 4 (https://creativecommons.org/licenses/by/3.0), alongside all other contributors and maintainers of the PhyloPic repository of free animal silhouettes. Finally, we would like to thank Cynthia Brewer for her ColorBrewer software (https://colorbrewer2.org), used in figure palettes.

