## Supplemental Materials for "Predator-like auditory stimuli elicit escape climb in *Aedes aegypti* mosquitoes"

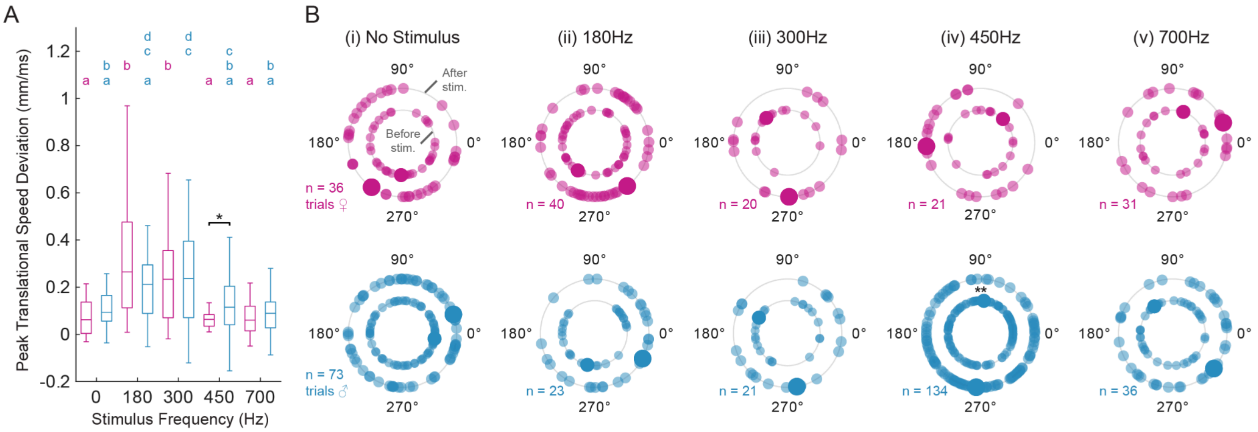


**Figure S1 Summary of translational speed and heading responses to auditory stimuli**

**(A)** Comparison of peak vertical speed from each trajectory (shown in B) between sexes and stimulus classes. Magenta and blue boxplots summarize female and male data respectively, with colored letters summarizing statistically distinct subsets within each sex (p< 0.05, One-way ANOVA familywise test with Tukey-Kramer *post-hoc* test). Black asterisk marks stimuli with statistically distinguishable differences between the two sexes (* = p < 0.05, two sample t-test), showing that males exhibit elevated vertical speed in response to the 450Hz female tone. **(B)** Summary of azimuthal heading over time for all free flight trajectories in response to each experimental tone. Stimulus speaker is located at the 0° azimuth. Small markers depict circular average heading for one trajectory during the 250ms preceding the stimulus (inner ring of markers) and circular average heading over the remainder of the video (outer ring of markers). Large markers depict the average of all trials. Black asterisks mark stimuli with statistically distinguishable distributions before and after stimulus onset, showing shift in azimuthal heading for male mosquitoes upon presentation of female wingbeat frequency tone (** = p<.01, Kuiper two-sample significant difference test for circularly distributed data).

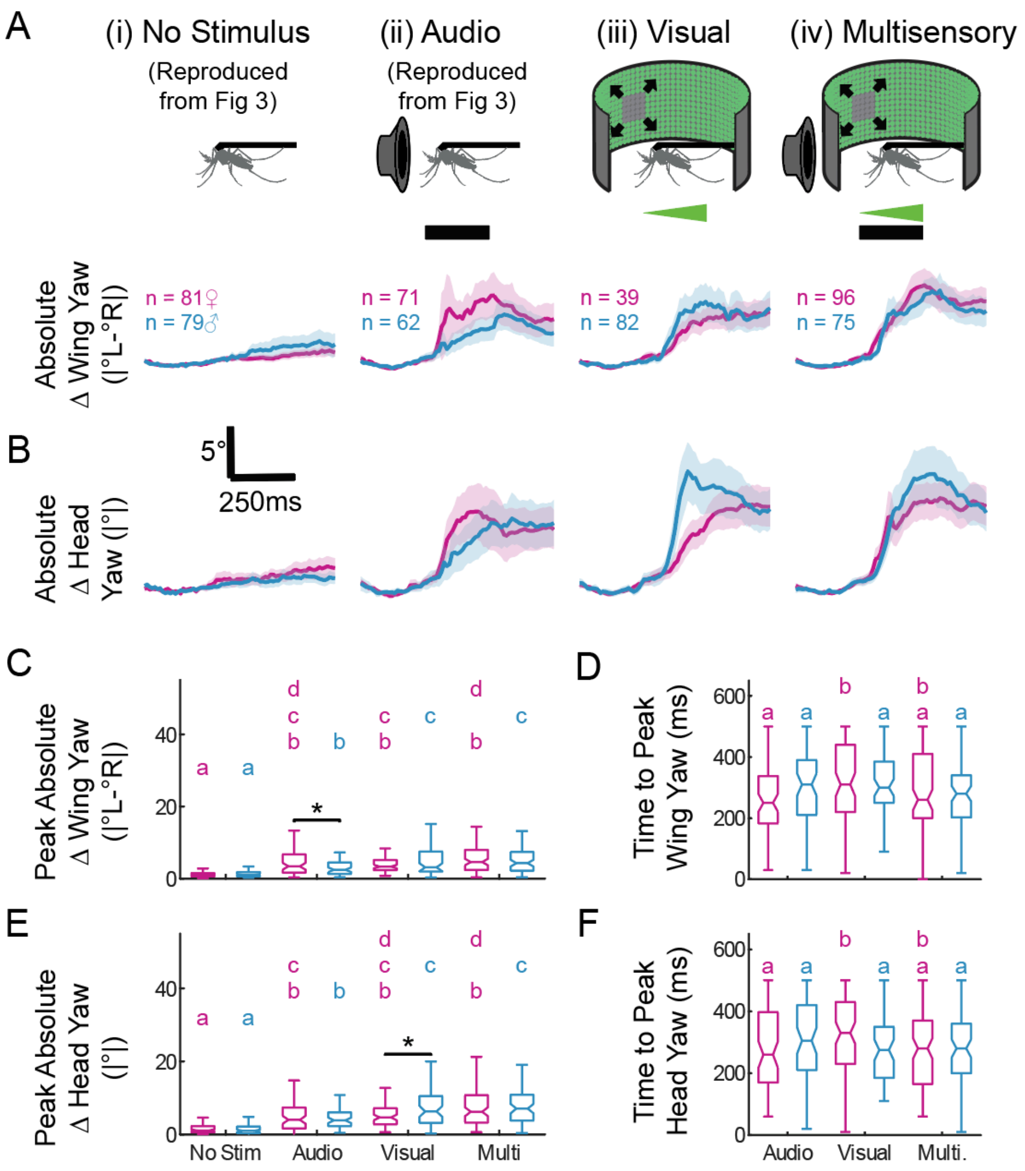


**Figure S2 Comparison of wing and head yaw during auditory, visual, and multisensory stimuli**
**(A)** Absolute wing yaw ($\Theta_L$-$\Theta_R$) change from pre-trial baseline over time, with schematic view illustrating each experimental condition: no stimulus (i) and 180Hz audio stimulus only (ii) reproduced from Figure 3 as well as (iii) presentation of an expanding square looming stimulus and (iv) concurrent presentation of both the looming and 180Hz auditory stimulus. **(B)** Same as (A), but instead showing absolute change from baseline for head yaw. **(C)** Comparison across sexes and conditions for absolute wing yaw response ($\Theta_L$-$\Theta_R$, colored letters reflect outcome of Kruskal-Wallis omnibus test with Conover-Iman *post-hoc* test within each condition, * = $p< 0.05$ Wilcoxon rank sum test pairwise between sexes), showing comparable response magnitudes for auditory, visual, and multisensory conditions for females. Response magnitude for visual and multisensory conditions exceeds those observed in response to unisensory auditory stimulation for males. **(D)** Same as (C), but for time to peak response across conditions, showing males and females responding at equivalent timescales under the visual and multisensory conditions, and

females responding more slowly to the visual stimulus than the auditory stimulus (** = $p < 0.01$ Wilcoxon rank sum test). **(E-F).** Same as C-D, but for absolute head yaw response.

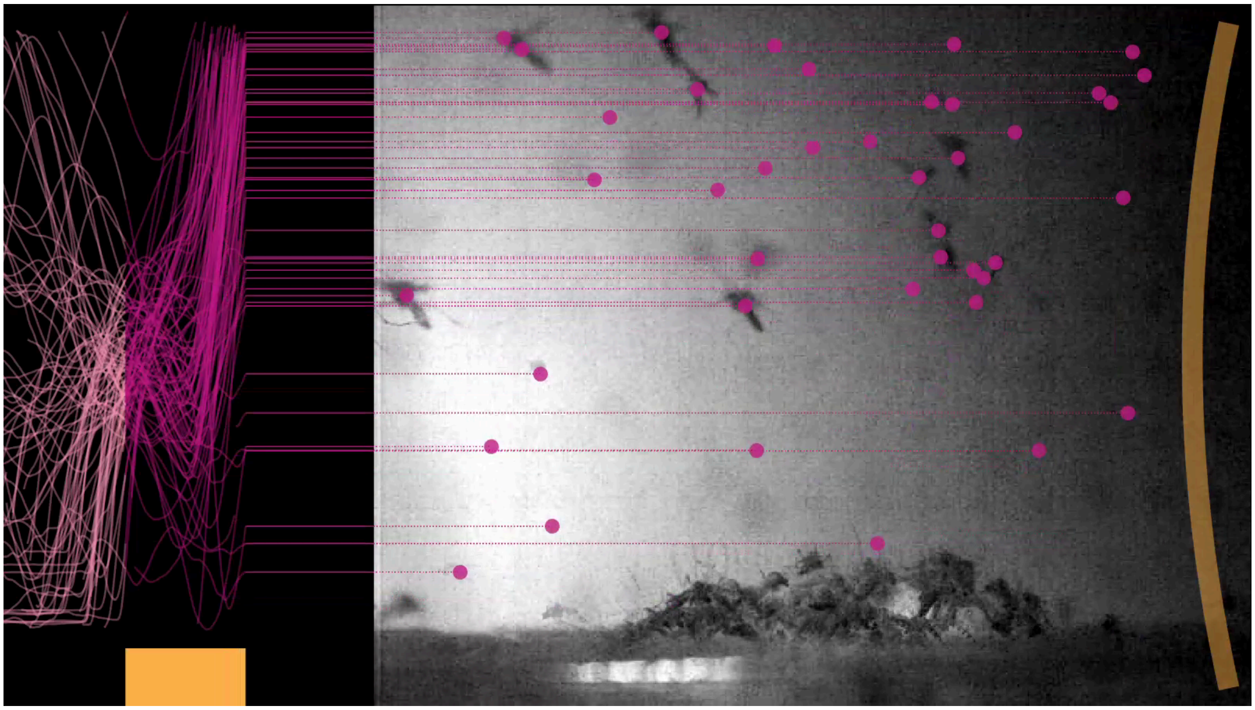

**Video S1 Female *Aedes aegypti* initiate rapid climb in response to 180Hz tone**
Video showing data from Figure 1C*ii*, aligning all videos and performing a minimum intensity projection on each time-aligned frame to visualize mosquitos from multiple videos. Vertical position over time for each trajectory is plotted in magenta, with stimulus epoch shown in yellow. Unlike data in Figure 1C, absolute position within video is shown rather than baseline-subtracted time series. Soundwaves originate from direction of the stimulus speaker, but do not illustrate individual cycles of stimulus waveform.

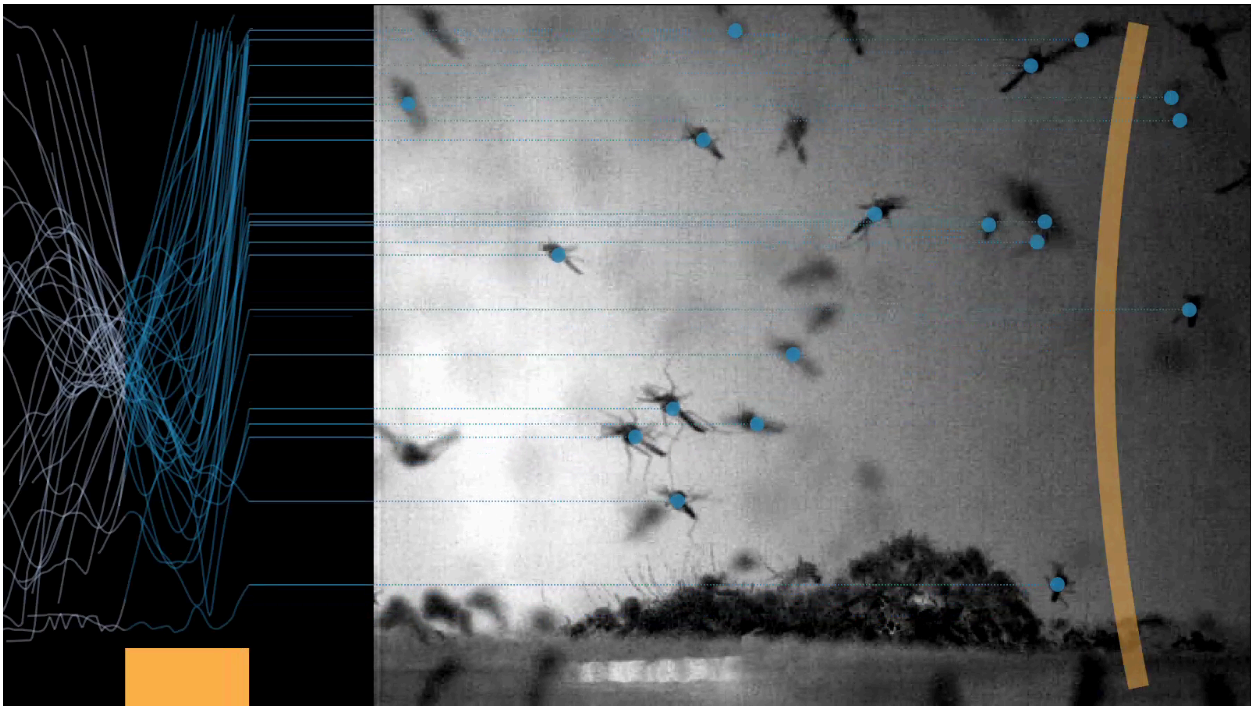

**Video S2 Male *Aedes aegypti* initiate rapid climb in response to 180Hz tone**
Same as Video S1, but instead showing the vertical position of male mosquitoes in blue.
